# Linear NeuroPaint: An Efficient Baseline for Cross-Session Neural Alignment and Inpainting

**DOI:** 10.64898/2026.10.03.756082

**Authors:** Yuxing Jared Yao, Ji Xia, Liam Paninski, Kenneth D. Miller

## Abstract

Large-scale Neuropixels recordings sample many brain areas, but no single experimental session records every area of interest. NeuroPaint, a transformer-based autoencoder, integrates such recordings by aligning area-specific latent dynamics across sessions and “inpainting” activity in unrecorded areas. Could a linear model, which would be computationally efficient and potentially more interpretable, suffice to achieve this? To address this question, we introduce **Linear NeuroPaint**, a linear variant of NeuroPaint, and evaluate it on two multi-area Neuropixels datasets. Linear NeuroPaint achieves strong predictive performance on recorded areas, however, it is less accurate than nonlinear models at cross-area inpainting, despite extensive hyperparameter optimization. Moreover, its parameters are not readily interpretable: incomplete cross-session alignment leads parameter-defined neuron clusters to exhibit heterogeneous response profiles. Together, these results show the limits of what a linear model can extract from multi-session recordings. Nevertheless, Linear NeuroPaint provides a computationally efficient baseline for benchmarking multi-session neural models.

## 1 Introduction

Neuropixels probes enable large-scale recordings across dozens of brain areas in behaving mice. However, no single experimental session records all areas of interest [7, 3, 1]. Integrating such recordings presents two related challenges. First, sessions that record the same brain area sample different neuronal populations, so their activity is expressed in different neural coordinate systems. These activity patterns must be aligned in a shared latent coordinate system to enable comparisons and information sharing across sessions [6, 10, 12, 2, 13]. Second, the aligned information must be combined across sessions to infer single-trial dynamics in brain areas that were not recorded in a given session, a problem we refer to as cross-area inpainting [11, 7].

NeuroPaint addresses both challenges with a transformer-based masked autoencoder [11]. It aligns area-specific latent dynamics across sessions and accurately predicts neural activity, including dynamics in brain areas not recorded in a given session. However, its nonlinear transformer backbone is computationally expensive and difficult to interpret. It remains unclear whether this nonlinearity is necessary: a carefully optimized linear model might perform comparably on some or all components of the problem. Distinguishing within-area prediction, cross-area inpainting, and cross-session alignment allows us to identify where nonlinear computation provides a meaningful advantage.

Previous studies have shown that linear transformations can align low-dimensional neural dynamics across recording sessions [4] and across animals [9]. These successes suggest that a linear model may be sufficient for cross-session alignment. However, these studies focused primarily on motor-related areas and did not address the joint problem of aligning sessions and across-area inpainting. Therefore, we introduce Linear NeuroPaint, a linearized variant that replaces NeuroPaint’s transformer backbone with a low-rank matrix. Its low computational cost permits extensive hyperparameter optimization, allowing us to establish a strong linear baseline and test which aspects of multi-session, multi-area modeling can be sufficiently captured by linear transformations.

Following [11], we evaluate our model on two multi-area Neuropixels datasets: the MAP dataset [3], and the IBL dataset [1]. On MAP, Linear NeuroPaint predicts neural activity in recorded areas comparably to LFADS [6], although both are outperformed by NeuroPaint. However, despite extensive hyperparameter optimization, Linear NeuroPaint is less accurate than both nonlinear models at cross-area inpainting. Its learned latent representations achieve only partial cross-session alignment, and its parameters are not readily interpretable: neurons grouped by similar model parameters can exhibit qualitatively different temporal responses. Ablation experiments nevertheless reveal that neural dynamics are broadly correlated across brain areas, with most areas contributing to a given target area. Together, Linear NeuroPaint provides a computationally efficient baseline for multi-session modeling, but is limited in what it can extract from the data for cross-area inpainting and cross-session alignment. Our results also demonstrate that model linearity does not necessarily confer interpretability.

## 2 Methods

### 2.1 Model

NeuroPaint is a masked autoencoder that infers dynamics in unrecorded brain areas by training across sessions with overlapping area coverage [11]. It consists of a cross-attention input stitcher, a transformer encoder, and a GLM readout stitcher with an exponential link function. The stitchers are session-specific, whereas the transformer is shared across sessions. In Linear NeuroPaint, we replace the cross-attention stitcher with a linear map and the transformer with a low-rank matrix (Fig. 1). The resulting model implements a low-rank linear mapping from unmasked neural activity to predicted log firing rates for masked data.

**Figure 1:**
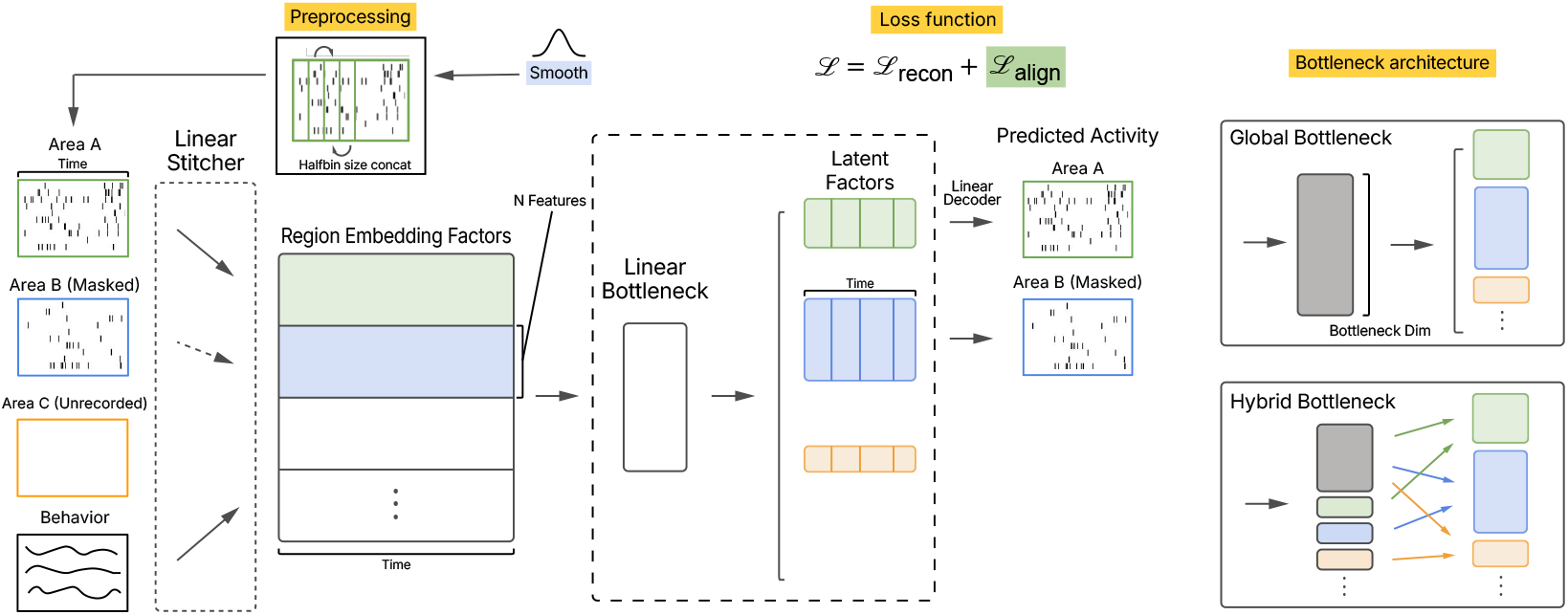
Linear NeuroPaint architecture. Linear NeuroPaint replaces the cross-attention stitcher and transformer encoder of NeuroPaint with linear maps, composing into a single low-rank operator. Two bottleneck variants are used: global (IBL) and hybrid (MAP).

We use 35 MAP sessions (aligned to sample onset, *−* 0.5 to 3.5 s) and 19 IBL sessions (aligned to stimulus onset, *−*0.5 to 1.5 s), with behavioral inputs of lick, jaw, and nose traces for MAP and pupil diameter, whisker motion energy, and wheel speed for IBL (Appendix A.6). We represent neural activity as spike counts in 10 ms bins and smooth the binned counts with a Gaussian kernel (*σ* = 5 bins; 50 ms). At each time point *t*, we concatenate the smoothed activity from bins *t −* 5 through *t* + 5, yielding an 11-bin (110 ms) temporal context window. NeuroPaint and LFADS see the full trial, but widening this window to 410 ms gave no clear gain (Appendix A.5). A session-specific linear stitcher projects the resulting features for each area into a region embedding. Hyperparameter optimization selected an embedding dimension of 48, a global bottleneck with hidden dimension 8 for IBL, and a hybrid bottleneck with 16 shared and 4 area-specific dimensions for MAP.

#### Training

The model is trained with one randomly selected area masked per training step (stitcher channels zeroed). It is trained to reconstruct neural activity of this area. Additionally, we randomly drop 20% of data in observed areas. We compute the reconstruction loss for all masked data with Poisson negative log-likelihood. To encourage cross-session alignment for inferred latent factors, we define a session-specific coding axis **a**_*s*_ for each of three task variables: stimulus, choice, outcome.

The alignment loss (ℒ_align_ = 1 − cos(**a**_*s*_, ā)) penalizes misalignment between each session’s coding axis **a**_*s*_ and a running cross-session consensus ā. The alignment loss is included with weight 0.05. Spike counts are used without centering, and behavioral inputs are z-scored per dimension within each session.

## 3 Results

### 3.1 Prediction performance

We evaluate prediction quality by DFE and *d*^*′*^(*t*) consistency (Appendix A.2 and A.3). On recorded areas, the linear model achieves DFE comparable to LFADS on MAP (0.152 vs. 0.149, Fig. 2A). All shared-latent-space models perform below the raw-spike baseline on recorded areas because this baseline reconstructs each neuron’s activity directly from its own spikes through Gaussian smoothing and the log-exponential transformation. It therefore serves as an empirical upper bound on prediction performance. The choice *d*^*′*^(*t*) discriminability trace computed from the model’s latent factors also correlates with *d*^*′*^(*t*) computed from smoothed spike data on recorded areas (median *r* = 0.64 on MAP, 0.37 on IBL, Fig. 2B; Appendix A.3).

**Figure 2:**
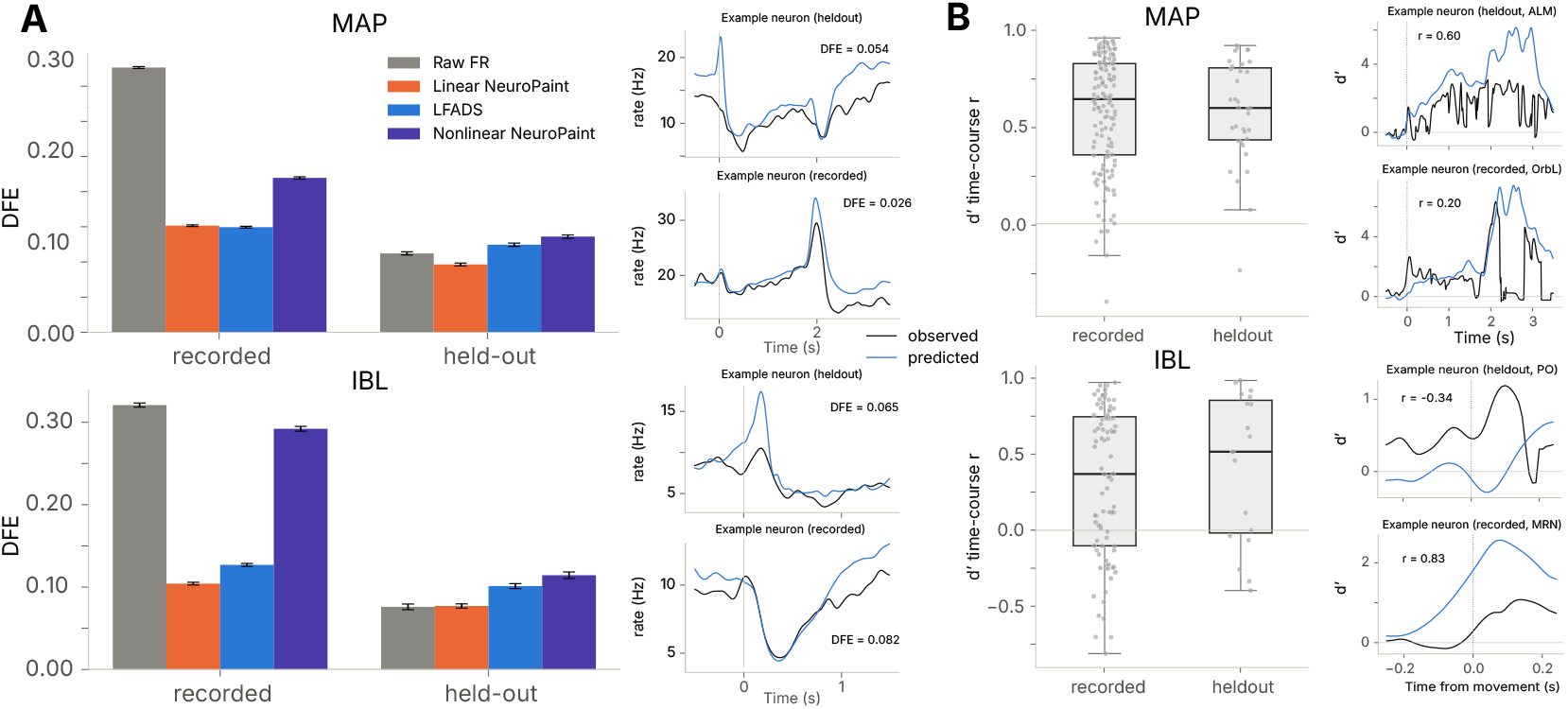
Prediction performance. **(A)** Spike-prediction DFE (*±* s.e.m.) for recorded and held-out areas, with example neurons’ observed (black) and predicted (blue) firing-rate traces. **(B)** Choice *d*^*′*^(*t*) time-course correlation (Pearson *r*) between inpainted and observed *d*^*′*^(*t*) across areas. A logistic regression decoder is trained to classify left-hit versus right-hit trials at each time bin, with its weight vector defining the decoding direction. At each time bin, *d*^*′*^(*t*) quantifies the separation between the two trial types after activity is projected onto this direction. We compute *d*^*′*^(*t*) from smoothed spike activity for the observed data (black) and from inferred latent factors for the predicted data (blue). (Appendix A.3). Linear NeuroPaint matches LFADS on recorded areas in MAP but otherwise underperforms nonlinear models, particularly on held-out areas.

Despite extensive hyperparameter optimization, the linear model does not match nonlinear NeuroPaint or LFADS on held-out areas (Fig. 2A). Its held-out DFE is comparable to that of the raw-firing-rate baseline, which is fit per session (0.096 vs. 0.112 on MAP, 0.077 vs. 0.076 on IBL). Nevertheless, the choice *d*^*′*^(*t*) trace from inpainted latents still correlates with that from raw firing rates on held-out areas (median *r* = 0.60 on MAP, 0.52 on IBL, Fig. 2B). The linear model thus recovers meaningful structure even where its DFE falls short.

### 3.2 Cross-session alignment

We assess alignment by cross-session decoding and trial-pair similarity (Appendix A.3). The linear model partially aligns neural activity across sessions. A logistic decoder for stimulus, choice, movement, and outcome trained on latent factors pooled across sessions retains 90–96% of the within-session ceiling (Fig. 3A). However, substantial session-to-session variability remains that linear alignment cannot explain away. Within-session trial-pair *r* averages 0.58, while across-session *r* averages only 0.23 (Fig. 3B). Per-dimension within-session *r* correlates with across-session *r* (Spearman *r* = 0.44, *p* = 4.9 *×* 10^*−*13^, Fig. 3C), showing that dimensions reliable within a session are also more consistent across sessions.

**Figure 3:**
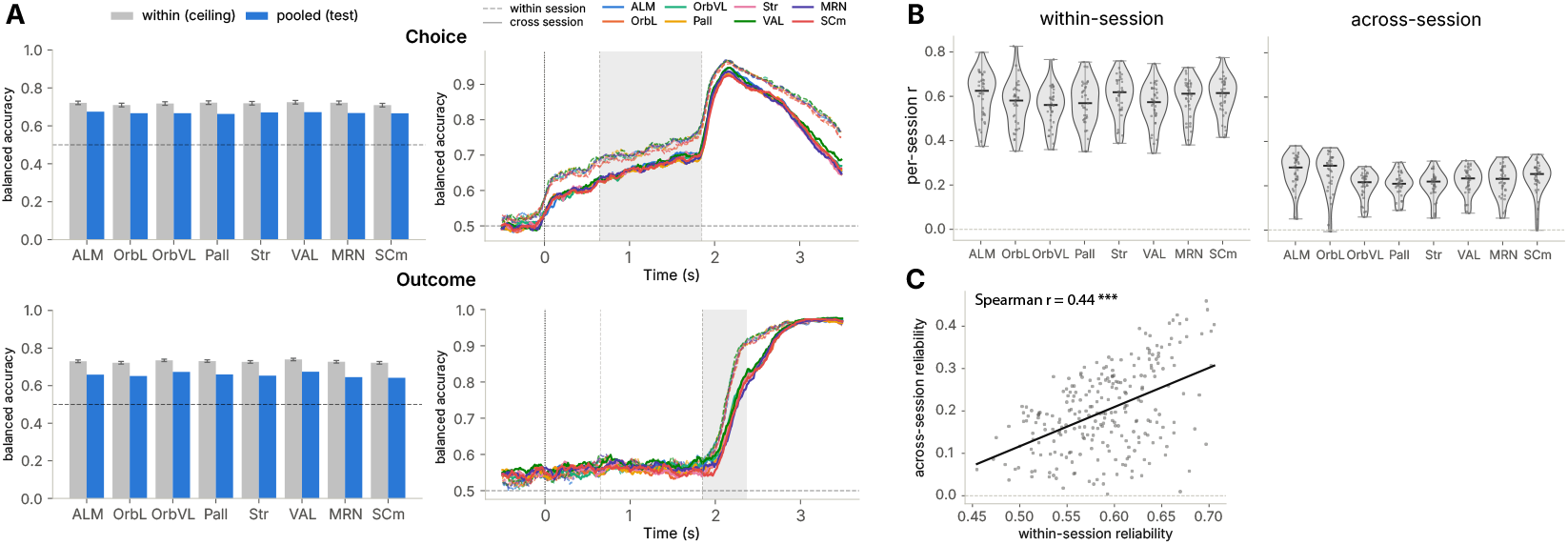
Cross-session alignment (MAP). **(A)** Logistic decoders for four task variables (stimulus, choice, movement, outcome). Bars show per-area balanced accuracy for within-session (gray) and cross-session (blue) decoders; shaded regions in time-course traces mark the scoring window. Choice (top) and outcome (bottom) displayed; see Fig. S4 for all variables. **(B)** Per-session trial-pair *r* for within-session (left) and across-session (right). **(C)** Per-latent-dimension within-vs. across-session reliability (MAP; Spearman *r* = 0.44, *p* = 4.9*×* 10^*−*13^). The pooled decoder retains 90–96% of within-session accuracy, but across-session trial similarity remains less than half of within-session similarity. See Fig. S5 for IBL.

### 3.3 Cross-session clusters of neurons based on model weights

We cluster neurons by their decoder-weight vectors and quantify task selectivity per cluster (Appendix A.3). The linear decoder gives each neuron an explicit weight vector onto the aligned latent space, making it natural to cluster neurons by their learned parameters rather than by trial-averaged PSTHs. We clustered neurons per area using KMeans with silhouette-based *k* selection (*k* = 3–20), removing single-session-dominated clusters [8]. In ALM, three clusters emerge (Fig. 4A), confirmed as statistically significant by a Gaussian null test. Although neurons within the same cluster have similar decoder weights, their PSTHs can differ (Fig. 4B) because the latent temporal profiles vary across sessions (Fig. 3B) and the model does not perfectly reproduce the observed firing rates (Fig. 2A).

**Figure 4:**
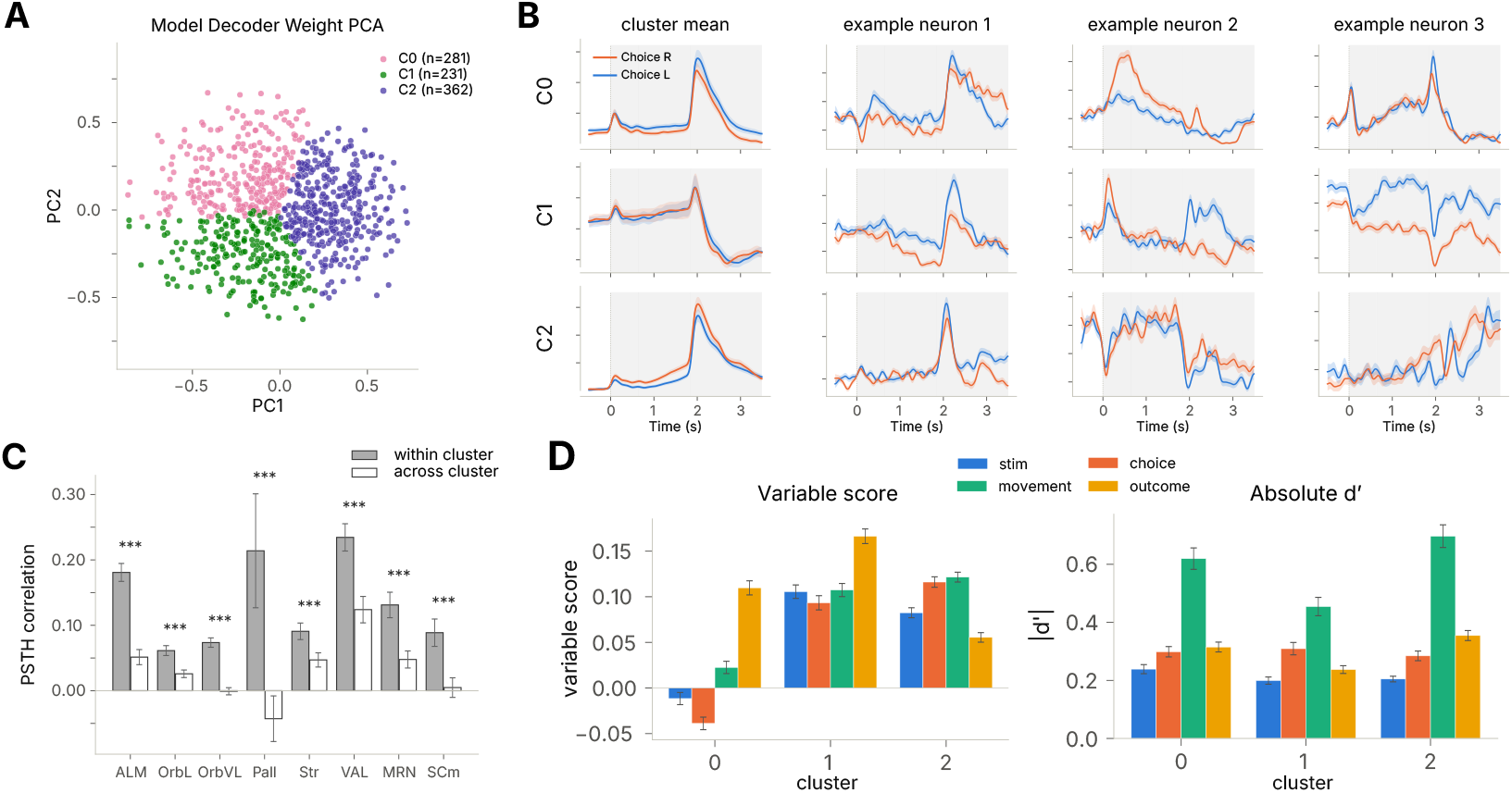
Cross-session clusters of neurons based on model weights (MAP, ALM). **(A)** PCA of per-neuron decoder-weight vectors, colored by KMeans cluster (*k* = 3; *n* = 281, 231, 362). **(B)**Per-cluster PSTH (choice L vs. R): cluster mean (left) and three centroid-nearest example neurons. **(C)**Within-vs. across-cluster PSTH correlation per area (permutation test, *p <* 0.001 all areas). **(D)**Per-cluster variable score (left) and raw|*d*^*′*^| (right) for stimulus, choice, movement, outcome; dashed lines show population mean. Clusters form statistically significant groups but contain neurons with heterogeneous temporal responses and mixed selectivity.

Within-cluster PSTH similarity is significantly higher than across-cluster similarity (*p <* 0.001 in all eight MAP areas, 2,000-shuffle permutation test, Fig. 4C), but the effect is modest, reflecting the limits of parameter-based clustering under session-to-session and trial-to-trial variability.

To further quantify what task-relevant information the clusters capture, we scored each neuron by the cosine similarity between its weight vector and the Haufe encoding pattern for each task variable [5]. We also computed per-neuron absolute *d*^*′*^ from raw firing rates to distinguish unselective neurons from those that are individually selective but whose preferred directions vary within the cluster, pulling the cluster-mean score toward zero. Cluster 0 scores near zero on stimulus and choice but has comparable raw |*d*^*′*^ |to the other clusters (Fig. 4D). Its neurons therefore have heterogeneous preferences for binary task variables rather than no tuning at all. Cluster 1 scores above the population baseline on all four variables, while cluster 2 shows clearer consensus on stimulus, choice, and movement but not outcome.

### 3.4 Area contribution

We quantify how much each source area contributes to predicting each target area by ablating its input and measuring the resulting drop in DFE (Appendix A.3, Fig. 5). The ablation matrix on recorded areas shows strong self-dependence: an area’s own input is the largest contributor to its own reconstruction, with diagonal DFE drops of 0.44–0.72 versus 0.08–0.29 off-diagonal in

**Figure 5:**
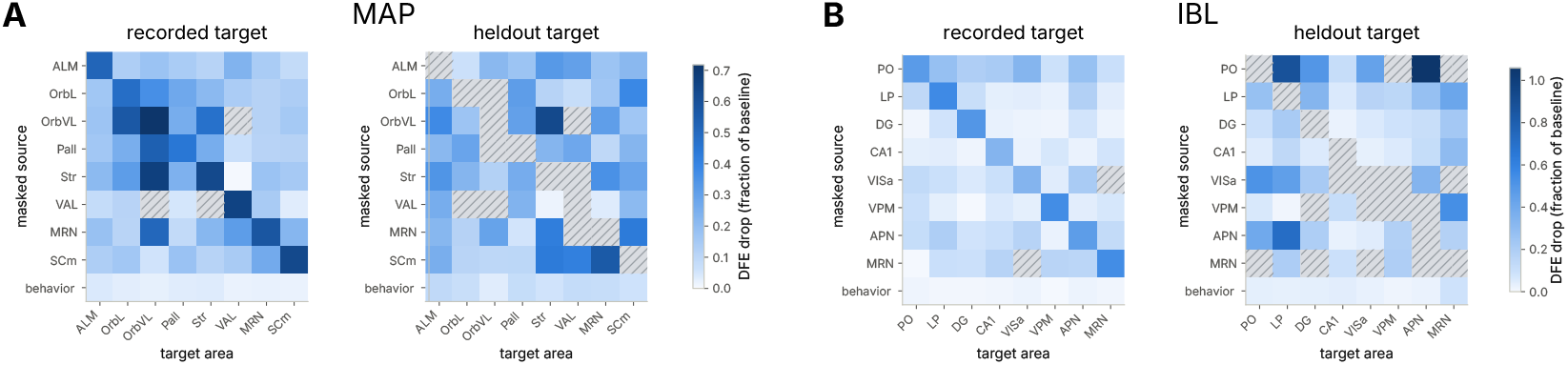
Area contribution. Fractional DFE drop when each source area is masked, gray cells indicate area pairs for which no data is available across sessions. **(A)** MAP: recorded (left) and held-out (right) output areas. **(B)** IBL, same layout. Self-dependence dominates the diagonal, but nearly every off-diagonal entry is positive, indicating broadly distributed cross-area contributions. MAP (Fig. 5A). Nearly every off-diagonal entry is positive. Neural dynamics are therefore broadly correlated across brain areas, and almost every recorded area contributes to explaining activity in a given target area. The held-out ablation matrices show similar cross-area interaction patterns (Fig. 5A–B, right), an independent validation on areas withheld from model training.

## 4 Discussion

Linear NeuroPaint shows that a shared linear transformation can partially align neural latent spaces across sessions but cannot accurately inpaint unrecorded brain areas. Across-session trial similarity remains less than half of within-session similarity, and this session-specific variability propagates into downstream analyses: neurons form statistically significant clusters in decoder-weight space, yet neurons within the same cluster can exhibit heterogeneous temporal responses. Ablation experiments reveal broadly distributed cross-area contributions. These results establish the linear model as a lower bound for cross-session neural inpainting and identify session-to-session variability as a central challenge for parameter-based neuron clustering. Future work should compare with a linear dynamical system (LDS) to disentangle the roles of temporal integration and nonlinearity, improve predictive accuracy, and develop statistical tests to assess the significance of entries in the area contribution matrix and their robustness to model degeneracy.

## A Appendix

### A.1 Model formulation and configuration

Figure S1 shows the full Linear NeuroPaint architecture. For each brain area *r*, raw spike trains 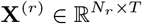 are smoothed with a Gaussian kernel (*σ* = 5 bins). At each time bin *t*, the smoothed activity from bins *t*−*h* through *t*+*h* (*h* = 5) is concatenated, yielding 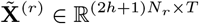.

A session- and region-specific linear stitcher maps the concatenated activity into a region embedding of dimension *D*:

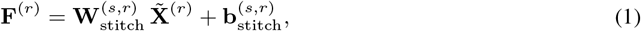

where 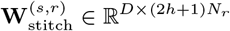 and 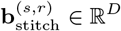 . The region embeddings for all *R* areas are stacked into a global feature matrix **F** ∈ ℝ^*RD×T*^, with zeros for masked or unrecorded areas.

A linear bottleneck compresses **F** to a low-dimensional representation. Three variants are used (Fig. S1). The **global** bottleneck projects all areas through a single shared map:

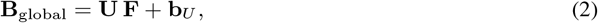

where **U** ∈ ℝ^*K×RD*^ and **B**_global_ ∈ℝ^*K×T*^ . Every area’s decoder receives the same bottleneck, so all cross-area information must pass through the *K*-dimensional shared subspace. The **area** bottleneck applies a separate map per area:

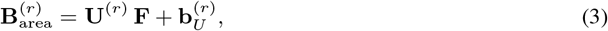

where 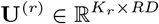. Each area has its own projection of the full region-embedding vector, so cross-area information is available but filtered through an area-specific lens. The **hybrid** bottleneck concatenates a shared projection with a per-area projection:

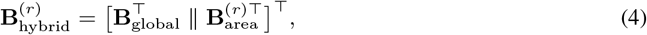

giving each area *K* + *K*_*r*_ latent dimensions that combine shared structure with area-specific variability. Hyper-parameter optimization selected *D* = 48 and *K* = 8 (global) for IBL, and *D* = 48 with *K* = 16 shared and *K*_*r*_ = 4 area-specific dimensions (hybrid) for MAP.

**Figure S1:**
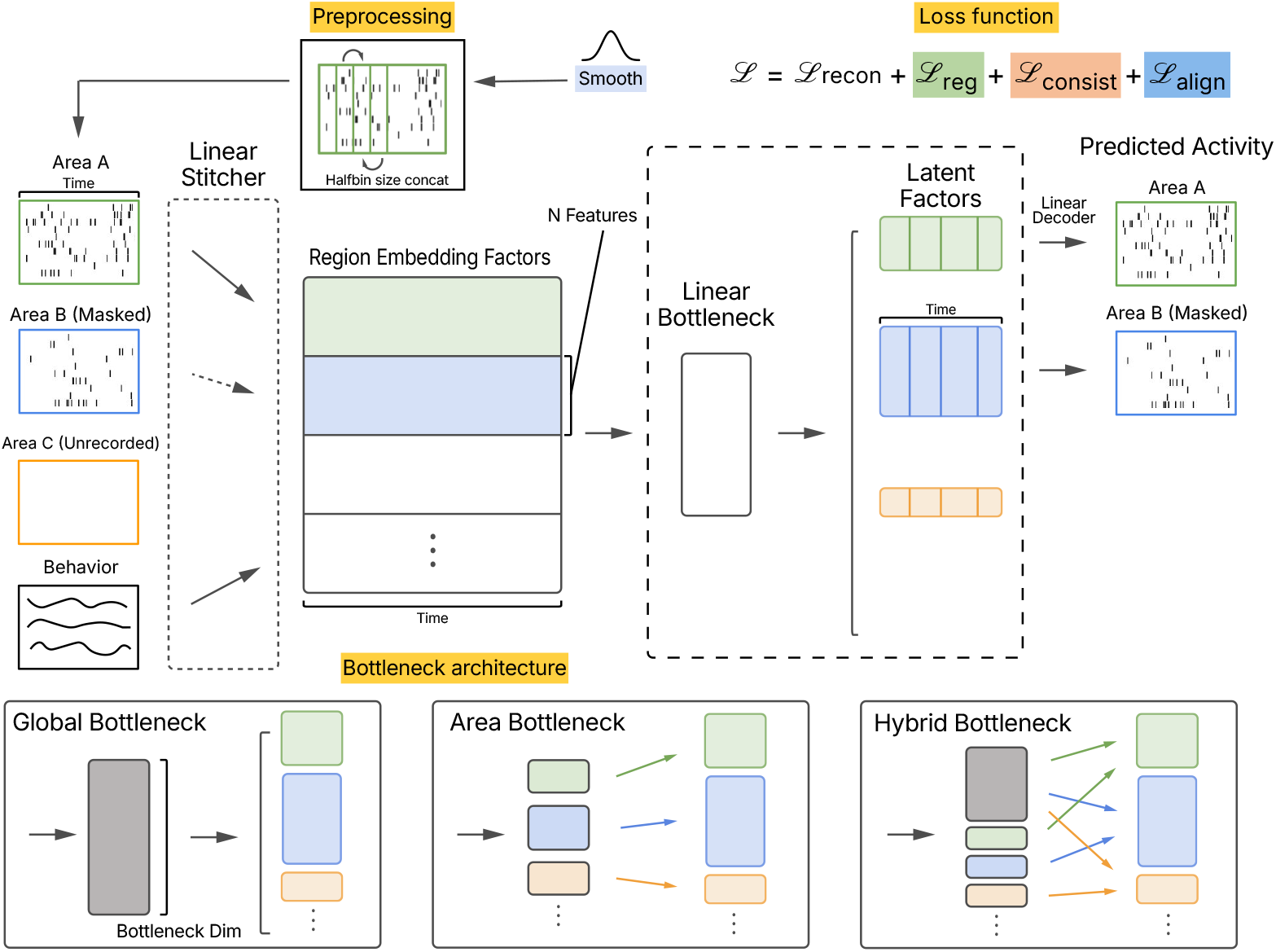
Full model diagram. Supplemental to Fig. 1. Linear NeuroPaint architecture showing preprocessing (Gaussian smoothing and sliding-window concatenation), per-session linear stitcher, region embedding, linear bottleneck, latent factors, and linear decoder. Bottom: the three bottleneck variants (global, area, hybrid). The loss function combines reconstruction loss ℒ_recon_ (Poisson NLL on masked area), regularization ℒ_reg_, consistency loss ℒ_consist_, and alignment loss ℒ_align_ (ℒ_reg_ and ℒ_consist_ have zero weight in the final models).

Region-specific decoders map the bottleneck to latent factors:

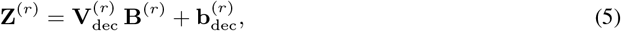

where **B**^(*r*)^ is whichever bottleneck variant is used, 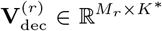 (*K*^*∗*^ being *K, K*_*r*_, or *K*+*K*_*r*_ respectively), and 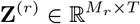.

Predicted firing rates are produced by a Poisson GLM readout:

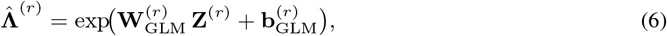

with 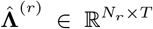. The total loss is ℒ = ℒ_recon_ + *λ* _reg_ ℒ_reg_ + *λ*_consist_ ℒ_consist_ + *λ*_align_ ℒ_align_, with *λ*_reg_ = *λ*_consist_ = 0 in the final models (Appendix A.4) and *λ*_align_ = 0.05. Here ℒ_consist_ is the consistency loss of Xia et al. [11], and the reconstruction loss is the Poisson negative log-likelihood

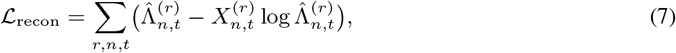

the regularization loss penalizes temporal roughness of the latent factors

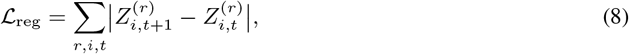

and ℒ_align_ is defined in Methods 2.

### A.2 Evaluation and baselines

#### Evaluation

Following Xia et al. [11], spike reconstruction is scored by DFE (fraction of Poisson deviance explained): a Poisson GLM is fit from the model’s latent factors to spikes per neuron, using separate train, validation, and test trial splits. One area per session is withheld from all model training (held-out); the remaining areas are the training set (recorded).

#### Baselines

We compare against two baselines evaluated with the same Poisson GLM: (1) LFADS [6] is a multisession variational sequential autoencoder with session-specific linear readin/readout layers, a shared GRU generator, and coordinated dropout. Posterior-sampled latent factors serve as the GLM predictors. (2) The raw-firing-rate baseline has no learned latent space: recorded spike trains are simply Gaussian-smoothed (*σ* = 5 bins) and log-transformed, then used directly as per-session GLM predictors.

### A.3 Analysis methods

#### Cross-session decoding

We decode four binary task variables: stimulus, choice, movement, and outcome (MAP) or reward (IBL). All decoders are fit on the training set and tested on the test set, using the same train/test trial split as model training. Each session’s latents are z-scored per dimension using training-trial statistics only. Training trials are capped at 50 for each ⟨stimulus, choice⟩ combination per session for class balance. For the pooled (cross-session) decoder, a logistic regression (*C* = 0.1, balanced class weights) is fit per time bin on training trials pooled across all sessions and evaluated on pooled test trials. The within-session decoder trains and tests within each session separately, serving as a ceiling since it does not require cross-session alignment. Balanced accuracy is reported over a task-relevant scoring window for each variable. On MAP (relative to sample onset): stimulus 0–0.65 s (sample epoch), choice 0.65–1.85 s (delay), movement 1.85–2.37 s (response), outcome 1.85–2.37 s (response). On IBL (relative to the variable’s anchor event): stimulus 0–200 ms post-stimulus onset, choice − 200–0 ms relative to movement onset (preparatory), movement 0–200 ms post-movement onset (execution), reward 0–200 ms post-feedback. The ratio of pooled to within-session accuracy quantifies alignment.

#### Cross-session similarity

For each pair of trials, we z-score each trial’s latent time course per dimension (excluding constant dimensions) and compute their Pearson correlation averaged over valid dimensions. Pairs are grouped into within-session (excluding self-pairs) and across-session bins. Separate trial types are computed independently and count-weight pooled, giving a per-session mean trial-pair *r* for each grouping. To assess per-dimension reliability, the same pairwise correlations are split into within- and across-session sums per latent dimension, and the mean within- and across-session *r* per dimension are compared with Spearman correlation.

#### Neuron clustering

Per area, each neuron’s decoder-weight vector is extracted from the linear decoder layer. Neurons with reconstruction loss above *µ* + 6*σ* of the bottom-half NLL distribution or mean firing rate below 1 Hz are excluded. KMeans is swept over *k* = 3–20 and the best *k* is chosen by silhouette score. After each sweep, clusters in which a single session contributes *>* 90% of the positive silhouette mass are removed and the sweep is repeated. To test whether the clustering captures structure beyond what a matched Gaussian would produce, we apply the null test introduced by Posani et al. [8]: 100 synthetic datasets are drawn from a Gaussian with the same mean and covariance as the real weight matrix, each is clustered with the same KMeans sweep, and the real silhouette score is compared against the null distribution (empirical *p*).

#### PSTH correlation test

To verify that decoder-weight clusters capture temporal response structure, we compute pairwise Pearson correlations of z-scored all-trial mean PSTHs for all kept neurons within each area. The test statistic is the mean within-cluster correlation minus the mean across-cluster correlation. Significance is assessed by a label-permutation test (2,000 shuffles). Standard errors are bootstrapped over neurons (1,000 resamples).

#### Cluster–variable score

Let **v**_*n*_ be the decoder-weight vector of neuron *n*. To obtain a latent direction associated with each task variable, we first balanced the trial set within each session (we select equal numbers of trials for each trial type), and then pooled balanced trials across sessions after the latent factors were *z*-scored separately within each session. For each task variable, at a given time bin *t*, a logistic discriminant weight **w**_*t*_ was mapped to Haufe pattern **d**_*t*_ = **Σ**_*t*_ **w**_*t*_ [5], where **Σ**_*t*_ is the covariance of the latent factors at the time bin *t* pooled across sessions. The per-bin patterns were averaged over the variable’s predefined task epoch to give an encoding direction **d**.

For each neuron, we defined variable score as the alignment between its decoder-weight vector **v**_*n*_ and the task variable’s encoding direction:

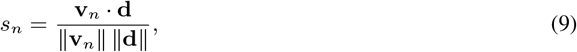

A positive score indicates that the neuron’s predicted firing rate is higher when the task variable takes the positive-reference value, i.e., right choice or stimulus. The cluster score was the mean of *s*_*n*_ over neurons assigned to that cluster.

#### Population discriminability *d*^*′*^(*t*)

For a binary task variable, a logistic decoder is fit at each time bin on the representation and trials are projected onto the decoder weight vector. The per-time-bin discriminability index is

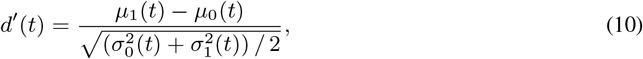

where *µ*_*c*_(*t*) and 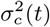 are the mean and variance of the projected scalar across trials in class *c* at time *t*. For inpainted *d*^*′*^(*t*), the representation is the model’s latent factors. For observed *d*^*′*^(*t*) (Fig. 2B), the representation is Gaussian-smoothed (*σ* = 5 bins) raw firing rates. To measure inpainting quality, we compute time-course consistency as the Pearson correlation between the *d*^*′*^(*t*) trace from inpainted latents and from raw recorded activity, evaluated on hit trials in the test split per masked area.

#### Single-neuron |*d*^*′*^|

Per-neuron discriminability |*d*^*′*^| applies the same formula to each neuron’s firing rate directly, without decoder projection. This measures each neuron’s single-unit selectivity independently of the model.

#### Area contribution

To measure each area’s contribution, each source area’s stitcher channels are replaced with zeros, and the DFE of every target area is recomputed with a frozen Poisson GLM (fit on baseline latents). The fractional DFE drop, (DFE_baseline_ − DFE_masked_)*/*DFE_baseline_, quantifies each source area’s causal contribution to reconstructing each target area.

### A.4 Loss configuration

We swept the weights of the two auxiliary losses from the original NeuroPaint paper (ℒ _reg_ and ℒ_consist_) across the three bottleneck variants (global, area, hybrid) on both datasets. The hybrid bottleneck with reconstruction loss only (ℒ_reg_ = ℒ_consist_ = 0) performed best on MAP, and the global bottleneck with reconstruction loss only performed best on IBL.

### A.5 Context window

NeuroPaint and LFADS access the full trial (4 s on MAP, 2 s on IBL) through self-attention and bidirectional recurrence, respectively. We swept the half-width *h* of the concatenation window (bins *t* − *h* through *t* + *h*) with the global bottleneck (*K* = 8) and input smoothing on. Table 1 reports held-out DFE. On MAP, increasing *h* from 5 to 20 improved DFE by only 0.0015. On IBL, performance peaked at *h* = 1 and decreased for larger windows, likely due to overfitting as the number of stitcher parameters grows with *h*. We therefore use *h* = 5.

**Figure S2:**
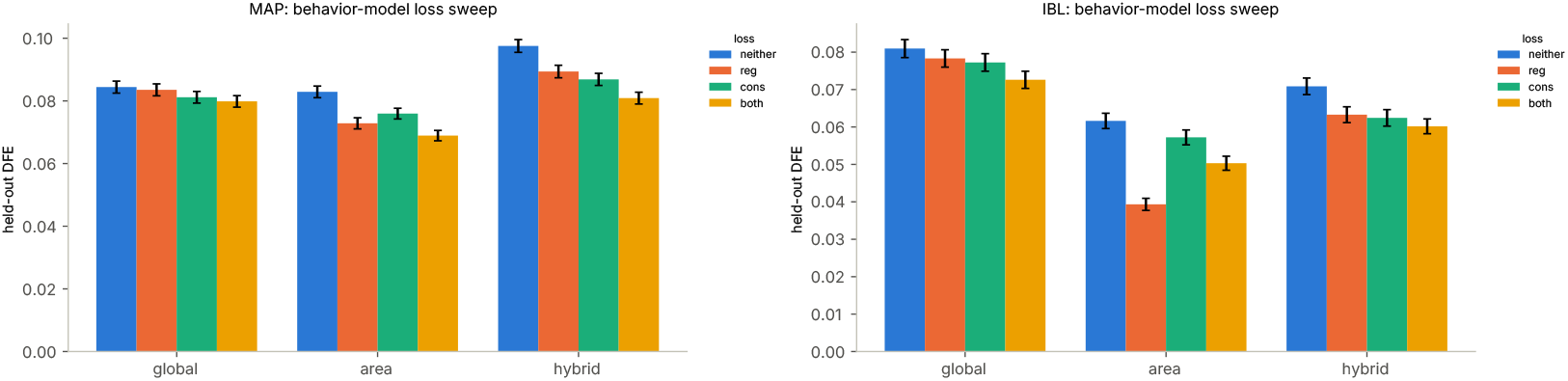
Loss weight sweep. Supplemental to Fig. 2. Held-out DFE across auxiliary loss weights (ℒ_reg_, ℒ_consist_) for the three bottleneck variants on both datasets.

**Table 1:** Context window sweep. Held-out DFE as a function of the window half-width *h* (worst 1% of neurons excluded). The window spans 2*h*+1 bins of 10 ms.

| *h* (window) | MAP | IBL |
|---|---|---|
| 0 (10 ms) | – | 0.0610 |
| 1 (30 ms) | – | 0.0654 |
| 3 (70 ms) | 0.0886 | 0.0636 |
| 5 (110 ms, used) | 0.0883 | 0.0641 |
| 10 (210 ms) | 0.0882 | 0.0584 |
| 15 (310 ms) | 0.0892 | 0.0526 |
| 20 (410 ms) | 0.0898 | 0.0603 |

### A.6 Behavior ablation

To assess whether behavior channels improve neural reconstruction, we compare the full model (neural + behavior inputs) against a model trained without behavior inputs. Including behavior consistently increases DFE across both datasets and area types (Fig. S3).

**Figure S3:**
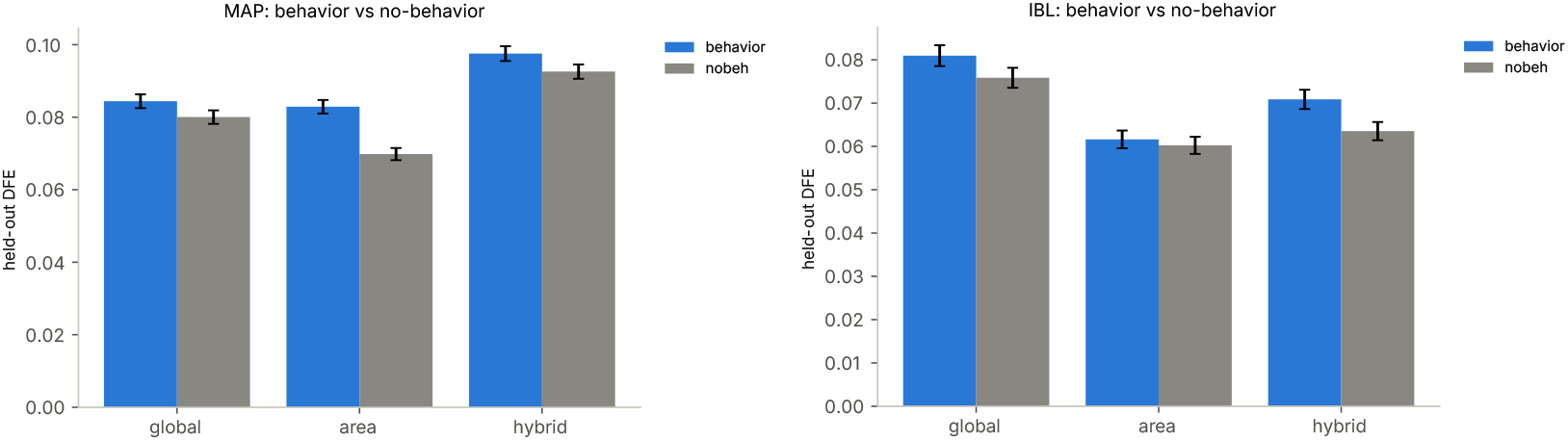
Behavior ablation. Supplemental to Fig. 2. Held-out and recorded DFE for the full model (neural + behavior inputs) versus the neural-only model on both datasets.

### A.7 Full cross-session decoding

Figure S4 shows within-session (ceiling) and cross-session (pooled) balanced accuracy for all task variables on both datasets. On MAP, the pooled decoder retains 90–96% of the within-session ceiling for stimulus, choice, movement, and outcome across all eight areas. On IBL, alignment ratios are similarly high (91–99%) for stimulus, choice, movement, and reward. Per-time-bin traces show that the pooled decoder closely tracks the within-session decoder throughout the trial.

**Figure S4:**
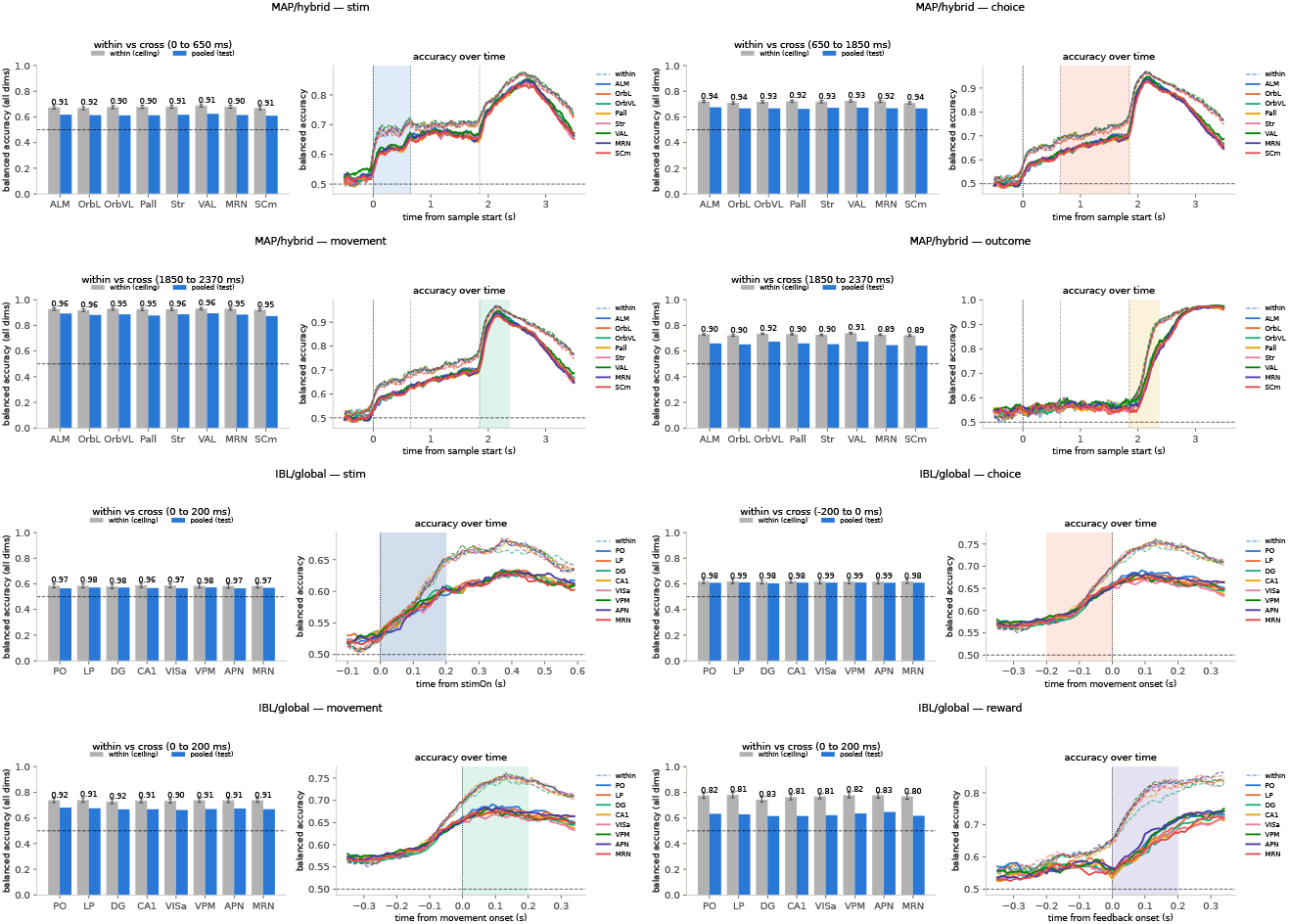
Full cross-session decoding. Supplemental to Fig. 3. Per-area balanced accuracy (bar, left) and per-time-bin traces (right) for within-session (ceiling) and cross-session (pooled) decoders. Top four rows: MAP/hybrid (stimulus, choice, movement, outcome). Bottom four rows: IBL/global (stimulus, choice, movement, reward). Numbers above bars are the pooled-to-within ratio.

### A.8 IBL alignment

Figure S5 shows the same session-similarity analysis for IBL. The same trend holds: within-session trial-pair *r* (∼ 0.35) is substantially higher than across-session *r* (∼ 0.12). Linear alignment leaves considerable session-to-session variability in this dataset as well. Unlike MAP, the per-dimension scatter shows no significant correlation between within- and across-session reliability (Spearman *r* = 0.03, *p* = 0.61), with some dimensions reaching across-session *r* above 0.15 while others fall near zero. This suggests that the global bottleneck produces a mixture of dimensions that generalize across sessions and dimensions that capture session-specific variance.

**Figure S5:**
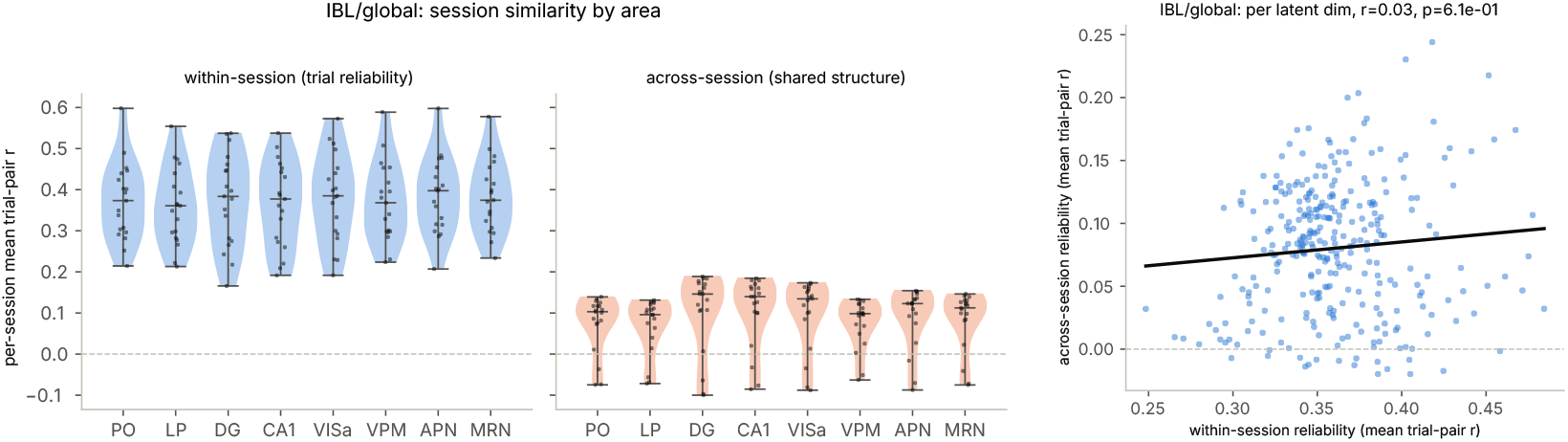
IBL cross-session alignment. Supplemental to Fig. 3. Trial-pair similarity and per-dimension reliability for the IBL dataset, same layout as Fig. 3B–C.

